# Simultaneous fMRI and fast-scan cyclic voltammetry bridges oxygenation and neurotransmitter dynamics across spatiotemporal scales

**DOI:** 10.1101/2021.05.28.446169

**Authors:** Lindsay R Walton, Matthew Verber, Sung-Ho Lee, Tzu-Hao Chao, R. Mark Wightman, Yen-Yu Ian Shih

**Affiliations:** Center for Animal MRI, University of North Carolina at Chapel Hill, Chapel Hill, NC USA; Biomedical Research Imaging Center, University of North Carolina at Chapel Hill, Chapel Hill, NC USA; Department of Neurology, University of North Carolina at Chapel Hill, Chapel Hill, NC USA; Department of Chemistry, University of North Carolina at Chapel Hill, Chapel Hill, NC USA

## Abstract

The vascular contributions of neurotransmitters to the hemodynamic response are gaining more attention in neuroimaging studies, as many neurotransmitters are vasomodulatory. To date, well-established electrochemical techniques that detect neurotransmission in high magnetic field environments are limited. Here, we propose an experimental setting enabling simultaneous fast-scan cyclic voltammetry (FSCV) and blood oxygenation-dependent functional magnetic imaging (BOLD fMRI) to measure both local tissue oxygen and dopamine responses, and global BOLD changes, respectively. By using MR-compatible materials and the proposed data acquisition schemes, FSCV detected physiological analyte concentrations with high spatiotemporal resolution inside of a 9.4 T MRI bore. We found that tissue oxygen and BOLD correlate strongly, and brain regions that encode dopamine amplitude differences can be identified via modeling simultaneously acquired dopamine FSCV and BOLD fMRI time-courses. This technique provides complementary neurochemical and hemodynamic information and expands the scope of studying the influence of local neurotransmitter release over the entire brain.

## Introduction

Coupling between neuronal activity and cerebral hemodynamic changes ensures that healthy energetic homeostasis is maintained within the active brain^1,2^. Classic blood oxygenation-level dependent functional magnetic resonance imaging (BOLD fMRI) data interpretation assumes that hemodynamic responses to stimuli are proportional to neuronal activity, such that hemodynamic response functions (HRF)^1^ can be derived to translate neuronal activity into BOLD signal predictions^3^. However, these assumptions are not always valid. Neurotransmitters have been receiving more attention as contributors to the hemodynamic response, as many are vasomodulatory (e.g., dopamine, nitric oxide, norepinephrine)^1,4^ and dysregulated in pathologies where neurovascular coupling is also dysregulated, such as schizophrenia and Parkinson’s^1,5^. It is paramount to understand how specific neurotransmitters influence vascular responses and accurately interpret neuroimaging data throughout the brain.

Dopamine is a vasomodulatory neurotransmitter concentrated in the striatum^4,6^, and plays a strong role in motivation, movement, addiction, learning, and reward prediction^7^. Dopamine receptors are located on microvasculature within the striatum, thalamus, and cortex, as well as on astrocytes^6,8^. Though dopamine has been studied intensely for decades, few studies have delved into its vascular modulation properties. Amphetamine and phencyclidine challenges show a linear relationship between evoked striatal hemodynamic changes and dopamine^9–11^, and D1- and D2-like dopamine receptor agonists show diametrically opposed vascular responses^6^. Though fMRI responses to cocaine and amphetamine challenge have been identified^10^, existing HRF models do not consider dopaminergic influence from shorter release events where higher affinity, presynaptic receptor binding would have more of an impact^9^. Dissecting how dopamine influences striatal vasculature is an important first step into understanding how whole-brain hemodynamics are affected by dopaminergic neurotransmission.

Popular methods of monitoring dopamine or other neurotransmitter kinetics *in vivo* are electrochemical, such as fast-scan cyclic voltammetry (FSCV) and amperometry^12^, though advances are being made with fluorescent sensor and magnetic resonance-based detection^13–16^, among others^17^. FSCV is especially desirable because it is chemically selective and quantifiable, and multiple, different neurotransmitters can be detected using well-established modifications^12,17^. It has high spatiotemporal resolution and minimal tissue damage, especially versus competing techniques like positron emission tomography and microdialysis^17^. FSCV translates directly to human use^18–20^, a challenge for fluorescent sensor-based techniques, and is therefore a promising method for studying neurotransmission during human behavior^19^, improving therapeutic deep-brain stimulation^21^, and interpreting human pharmacological fMRI studies^22^. Though FSCV can detect both tonic^23^ and phasic dopamine changes, and oxygen changes^24–27^, its high spatial resolution limits the scope of interpretation. To reveal the influence of local dopamine release on whole brain circuits requires pairing FSCV with fMRI to acquire neurotransmitter and brain-wide hemodynamic data simultaneously.

In this work, we develop and characterize a simultaneous FSCV-fMRI technique. We address the challenges of developing an MR-compatible FSCV recording system, circumventing high-frequency electronic noise from MR imaging gradients, and synchronizing multimodal data acquisition. We perform *in vivo* FSCV-fMRI during an oxygen inhalation challenge using an oxygen-sensitive FSCV waveform and electrical deep brain stimulation using both oxygen- and dopamine-sensitive FSCV waveforms at an electrode implanted into the nucleus accumbens (NAc). To compare hemodynamic measurements at different spatial scales, we collected tissue oxygen and BOLD fMRI data concurrently. We derive an HRF from high-resolution dopamine data acquired during simultaneous fMRI, and for the first time, we demonstrate that this simultaneous FSCV-fMRI recording platform can identify brain regions that encode dopamine amplitude changes. This method should contribute significantly to the understanding of local neurotransmission on dynamic changes of brain activity.

## Results

### fMRI and FSCV compatibility

A major step to achieve simultaneous FSCV-fMRI is ensuring that FSCV material components do not produce imaging artifacts. Traditional glass capillary microelectrodes, like those used for FSCV, produce minor artifacts^28^, but polyimide-fused silica further enhances MR-compatibility^29^ and is also used with FSCV microelectrodes (Fig.1A)^30^. Both in agarose phantoms and *in vivo*, polyimide-fused silica electrodes showed minimal MR artifacts (Fig.1A-C); however, modifications were necessary to maximize MR compatibility. Standard fabrication connects the carbon fiber directly to a connection pin via silver epoxy, which provides structural support within the headcap and makes the electrode suitable for chronic implantation^30^. Unfortunately, commercially available pins contain trace amounts of nickel, a magnetic material with large susceptibility artifacts^31^. Here, we silver epoxied the carbon fiber directly onto silver wires. Following implantation, 1 cm of silver wire from both working and Ag/AgCl reference electrodes was left exposed above the headcap during recovery (Fig.1D). To protect the free-standing wires, plastic shields were cemented to the headcaps (Fig.1D). We performed flow-through analysis to assess whether these modifications affected sensitivity. In our hands, the calibration factors for oxygen and dopamine on the oxygen-sensitive waveform were −0.19 nA/µM/100 µm and 4.8 nA/µM/100 µm, respectively (Supplemental Fig.1). Both calibration factors fall between values reported for glass microelectrodes (normalized to 100 µm)^24^. On the dopamine-sensitive waveform, dopamine sensitivity was 34 nA/µM/100 µm, in agreement with the literature^30^ (Supplemental Fig.1). These data indicate that our modified polyimide-fused silica electrodes are MR compatible and can detect analytes of interest with high sensitivity.

**Fig. 1.**
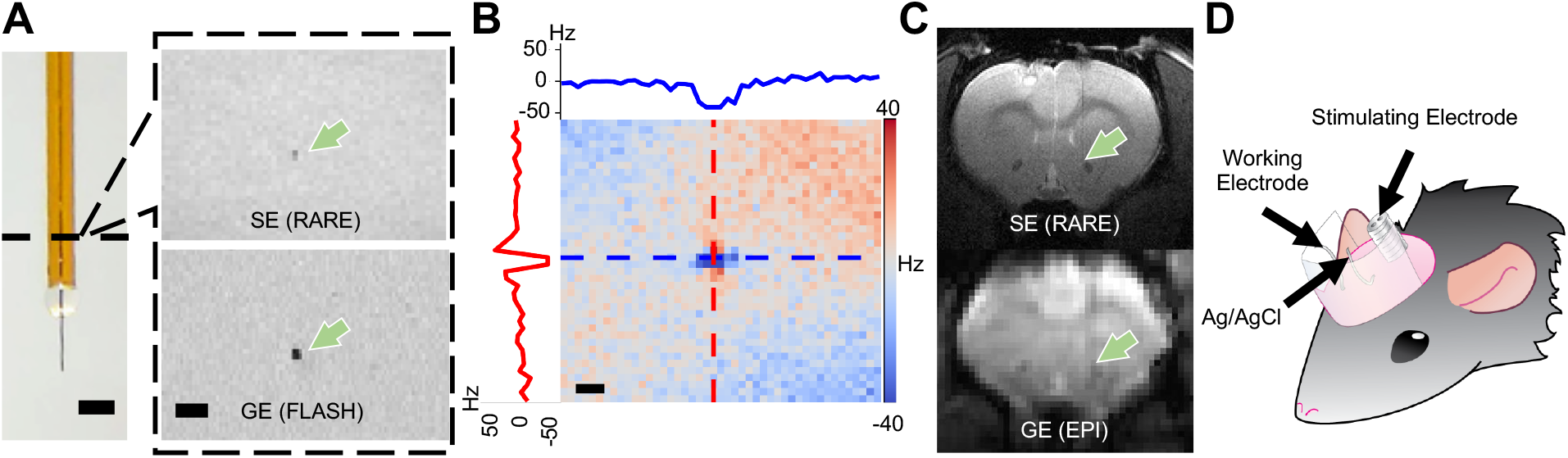
Modifying FSCV materials improves MRI compatibility. (A) *Left*: Polyimide-fused silica capillary microelectrode (scale bar=100 µm). *Right*: Electrode axial cross-section scanned inside an agarose phantom using spin echo (SE) rapid acquisition with relaxation enhancement (RARE), and gradient echo (GE) fast low angle shot (FLASH) sequences (scale bar=1 mm). (B) Axial view of the magnetic field map showing inhomogeneities induced by the microelectrode materials in an agar phantom. Scale bar=200 µm. Frequency profiles are taken from the dashed line cross-section of the same color (horizontal=blue, vertical=red). (C) Microelectrodes implanted in the rat NAc *in vivo*, indicated with arrows, during a SE-based RARE anatomical scan and a GE-based echo planar imaging (EPI) functional scan. (D) Rat headcaps were fitted with plastic shields to protect loose silver wire connections to FSCV working and reference electrodes.

FSCV is a differential technique that detects electrooxidation and reduction reactions on the scale of nA or less (Supplementary Fig.2), so electrical noise must be minimized to maximize FSCV sensitivity. First, we placed a dummy cell (*vide infra*) inside the MRI bore to quantify ambient electrical noise. Sub-nA noise was achieved using minimal grounding and braided cables with the gradients disabled, but a 4^th^ order Bessel low-pass filter with a 2 kHz cut-off frequency was necessary to remove the high amplitude, 50 kHz power amplifier noise present with enabled gradients (Supplementary Fig.3). To ensure that noise levels were low enough to detect physiologically relevant concentrations of analyte, we performed a flow-injection experiment using dopamine boluses. A flow-injection cell was placed inside the MRI bore on a 3D printed holder attached to the animal sled (Fig.2A). With repeated boluses, there was a significant main effect of the recording environment on the signal-noise-ratio (SNR; F_(1.8, 5.3)_=40.9, p=0.0007, Fig.2B). Boluses recorded with the gradient enabled and without low-pass filtering had significantly lower SNR (6.46±2.63) than those recorded with low-pass filtering (disabled with filter: 135.9±13.9, enabled with filter: 89.5±16.0; Tukey’s post-hoc, p=0.003 and p=0.03, respectively; Fig.2B). Our chosen filter did not distort bolus amplitudes or shapes (Fig.2C-D), making it an effective method of removing the majority of electrical noise within the MRI bore.

**Fig. 2.**
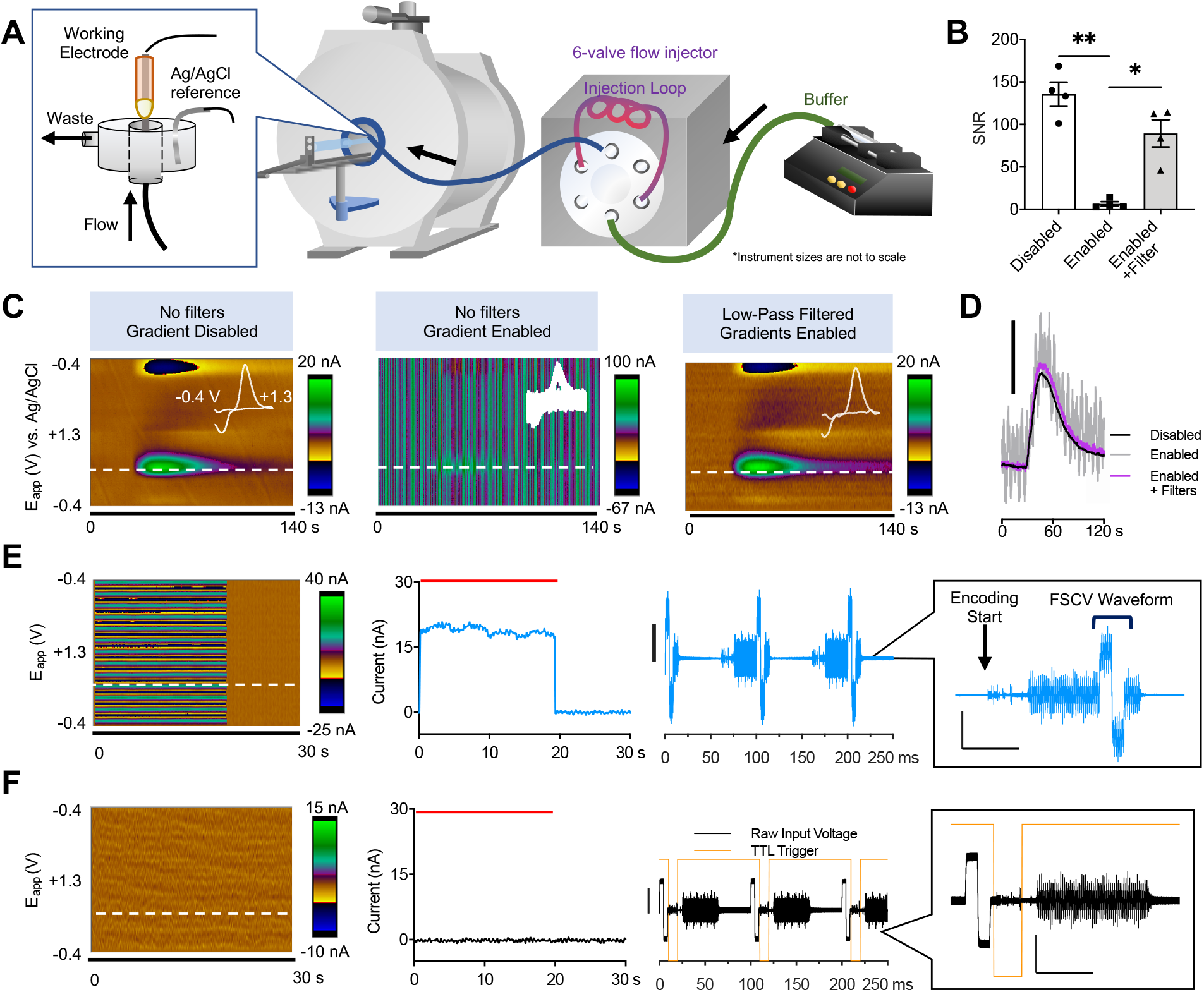
Software and hardware modifications for simultaneous FSCV-fMRI. (A) Within-bore flow-injection analysis requires the 6-port injection valve to be placed outside of the magnet at a safe distance. A fresh dopamine bolus is injected and flowed through to the electrode inside of the MRI bore for detection. (B) SNR for a physiologically relevant dopamine concentration. Averaged triplicate measures are shown (n=4 electrodes; Tukey’s post-hoc *p<0.05, **p<0.01). (C) Low-pass digital Bessel filters remove the high-frequency noise from enabled MRI gradients without distorting FSCV signals. Color plots indicate flow-injection dopamine boluses within the MRI bore. Cyclic voltammograms obtained from the maximum bolus current are overlaid in white. (D) Dopamine bolus injection time-courses from the dopamine oxidation potential in C. Scale bar=15 nA. (E) Dummy cell data collected inside the MRI bore. *From left to right:* Color plots, current time-courses acquired at 10 Hz, and raw voltage time-courses acquired at 100 kHz (scale bar= 4V) with corresponding magnification insets (vertical scale bar=1 V, horizontal scale bar=20 ms). Large artifacts appear when FSCV waveform applications and EPI encoding overlap temporally, but (F) interleaving the signals with TTL triggers avoids this interference. Time-courses are taken from the dopamine oxidation potential, indicated by a white dashed line in the color plots. Red bars indicate EPI scan duration.

### Simultaneous fMRI and FSCV

It is well-documented with other electrical recoding modalities (e.g., electrophysiology) that echo planar imaging (EPI)-based fMRI produces intrusive, high-amplitude noise during encoding, which can be removed via principal component regression, algorithms, data acquisition interleaving, or deleting encoding portions outright^3,32–34^. FSCV detects electrochemically active analytes via periodic electrooxidation/reduction voltage sweeps (Supplementary Fig.2). As EPI-based fMRI also delivers RF and gradient pulses periodically, ideally FSCV waveforms could interleave with active RF and gradient pulsing. A dummy cell was placed inside the bore during fMRI acquisition to observe the encoding artifact, which was approximately 2 V_P-P_, 55 ms, and repeated with each slice acquisition (Fig.2E-F). Any temporal overlap between the waveform application and EPI encoding resulted in high amplitude interference (Fig.2E). To interleave FSCV and EPI recordings, MRI protocols were set to wait for a TTL trigger per slice and all timings were coordinated within the FSCV data acquisition software. A 10-slice EPI scan using a 1000 ms repetition time (TR; i.e., 100 ms between EPI artifacts) could interleave with FSCV waveforms applied at 10 Hz (i.e., 100 ms between waveform scans) for 10:1 FSCV:fMRI temporal resolution acquisition. When FSCV data acquisition begins, the software sends a high TTL voltage to the MRI. After each FSCV waveform is applied, the TTL remains high for an additional 3 ms before a 10 ms, low voltage TTL triggers EPI slice acquisition (Fig.2F). By using TTL trigger-dependent interleaving, one or more FSCV scans can be collected per EPI slice without encoding gradient interference.

### In Vivo Oxygen Challenge in the NAc

Previous FSCV studies have shown that pharmacologically manipulating neurotransmission affects oxygen dynamics, which are taken to represent hemodynamic changes^24,25,27^; however, the spatial resolution of FSCV limits the interpretation to highly localized microenvironments. Using an oxygen-sensitive waveform (Fig.3A), we simultaneously acquired oxygen-related signals from FSCV and fMRI, *in vivo*, at spatial scales of different magnitudes (Fig.3B). We first administered a 100% oxygen breathing challenge (Fig.3A) and compared evoked responses within subject-level regions of interest (ROI) at the electrode tip implanted in the NAc (Supplementary Fig.4). Both fMRI and FSCV oxygen changes increased during hyperoxia and fell to baseline afterwards (Fig.3C, n=6 subjects). The 40 s delay before oxygen levels change corresponds to the transit time between the gas flow source and the ventilated subject. FSCV data, binned to match fMRI temporal resolution (TR=1000 ms), had significantly higher SNR during hyperoxia (FSCV: 47.7±13.8, fMRI: 4.4±0.4; t_(17)_=8.24, p<0.0001, n=18 events; Fig.3D). The averaged event time-courses highly correlated (r=0.85, p<0.0001, n=18 events; Fig.3E) in agreement with the literature^35^; however, there were differences between the observed oxygen kinetics (Fig.3F, n=18 events). The FSCV response preceded the highest correlated BOLD responses by 5±2 s, according to time shift analysis (n=18 events; Fig.3F). The clearance time from the maximum response to half-max was significantly longer in BOLD fMRI versus oxygen FSCV (n=6 subjects, FSCV: 20.7±3.0 s, fMRI: 33.7±3.4 s; t_(5)_=4.48, p=0.007; Fig.3G), but there was no difference in half-max rise times. The data trended towards a longer fMRI versus FSCV full-width at half-max (FWHM; n=6 subjects, t_(5)_=2.21, p=0.08; Fig.3G), but did not reach statistical significance. Though FSCV and fMRI detect highly correlated changes during an oxygen inhalation challenge, the difference in clearance kinetics indicate that these data are complementary. As expected of a systemic, uniform increase in oxygen from an inhalation challenge, induced BOLD responses and FSCV oxygen highly correlated spatially across the brain (one sample t-test, p=0.05 compared to other voxels, followed by group statistics with n=6 subjects; Fig.3H). This analysis demonstrates the feasibility of using local FSCV signals for correlation/regression analysis at a whole brain scale.

**Fig. 3.**
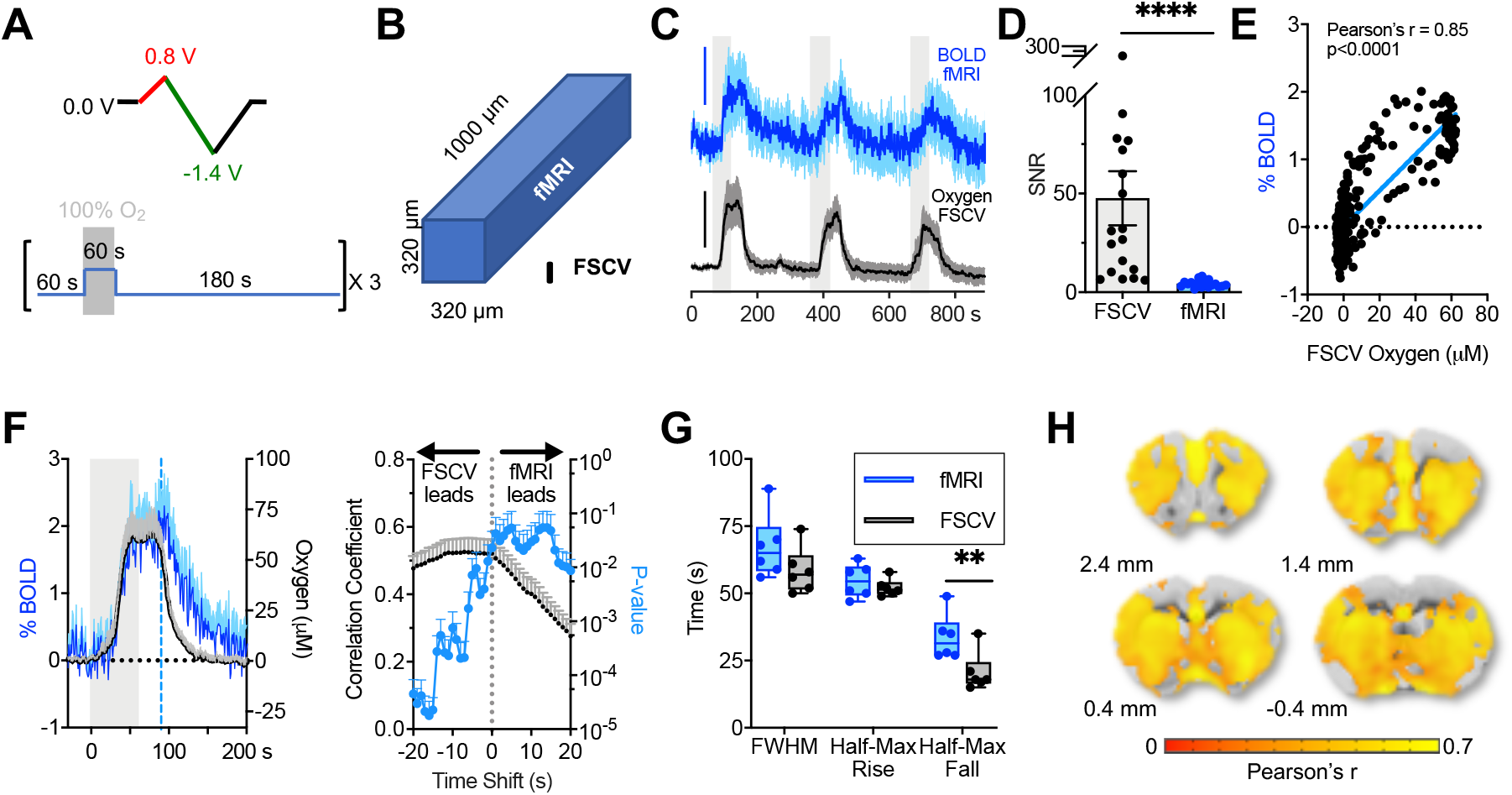
BOLD fMRI and oxygen FSCV time-courses in the rat NAc highly correlate during oxygen breathing challenges. (A) Paradigm schematic. *Top*: An oxygen-sensitive waveform was repeated throughout the recording session to detect oxygen changes. Red indicates the oxidation potential sweep, green indicates the reduction potential sweep. *Bottom:* Oxygen inhalation event paradigm. When not inspiring pure oxygen, subjects inspired medical air. (B) A single BOLD fMRI voxel compared to the volume sampled by FSCV, to scale. (C) BOLD fMRI and oxygen FSCV time-courses show signal increases during oxygen inhalation periods (n=6 subjects). Blue scale bar=2% signal change. Black scale bar=50 µM oxygen. (D) SNR from FSCV and fMRI NAc ROI oxygen inhalation time-courses (n=18 events, ROIs=36±3 voxels). (E) Averaged BOLD fMRI and oxygen FSCV time-course correlation (n=18 events). (F) Averaged oxygen challenge events from C (*left*) and corresponding time shift-correlation plot (*right*). (G) Kinetic parameters derived from subject-averaged time-courses, where half-max rise is the time from t=0 to half-max, and half-max fall is the time from t=30 s after the 100% oxygen ends (indicated in f as a blue dashed line) to the half-max clearance value (n=6 subjects). (H) Group-level correlation between BOLD fMRI and concurrently acquired oxygen FSCV time-courses (p=0.05). Voxels with correlation coefficient <0.3 are excluded. Grey boxes in A, C, and F indicate times where gas inhalation lines were switched to 100% oxygen. Error bars are ±SEM or +SEM, for visual clarity. Ratio-paired t-test **p<0.01, ****p<0.001.

### In Vivo Electrical VTA Stimulation

BOLD fMRI and oxygen FSCV map stimulus-evoked responses in the brain on different scales^2,24,25^. To demonstrate how FSCV-fMRI can study specific stimulus-evoked responses, we electrically stimulated the ventral tegmental area (VTA; Fig.4A). Evoked responses had higher SNR with FSCV versus fMRI responses within the NAc ROI (FSCV: 17.4±3.2, fMRI: 3.3±0.6; t_(17)_=5.40, p<0.0001, n=18 events; Fig.4B). Stimulations evoked positive oxygen FSCV and BOLD fMRI changes, which are characteristic of electrical VTA stimulation^24^ (Fig.4C). The averaged time-courses highly correlated (r=0.83, p<0.0001, n=18 events; Fig.4D). FSCV time-courses lagged the maximally correlated fMRI time-courses by 0.8±0.8 s, according to time shift analysis (Fig.4E). While traditional block-design general linear model (GLM) analysis extracted broad areas of VTA-induced BOLD responses (Supplementary Fig.5), similar to those reported in the literature^7,36,37^, the brain areas showing the highest oxygen FSCV time-course correlations were more tightly localized to clusters in ipsilateral NAc, claustrum, and insular cortex, bilateral prelimbic and cingulate cortices, and septum (one sample t-test cutoff at p=0.05 compared to other voxels in the brain, followed by group GLM analysis with n=6 subjects; Fig.4F). FSCV oxygen measurements could be used to identify subsets of traditional GLM-identified voxels that most closely match the hemodynamic response of an ROI.

**Fig. 4.**
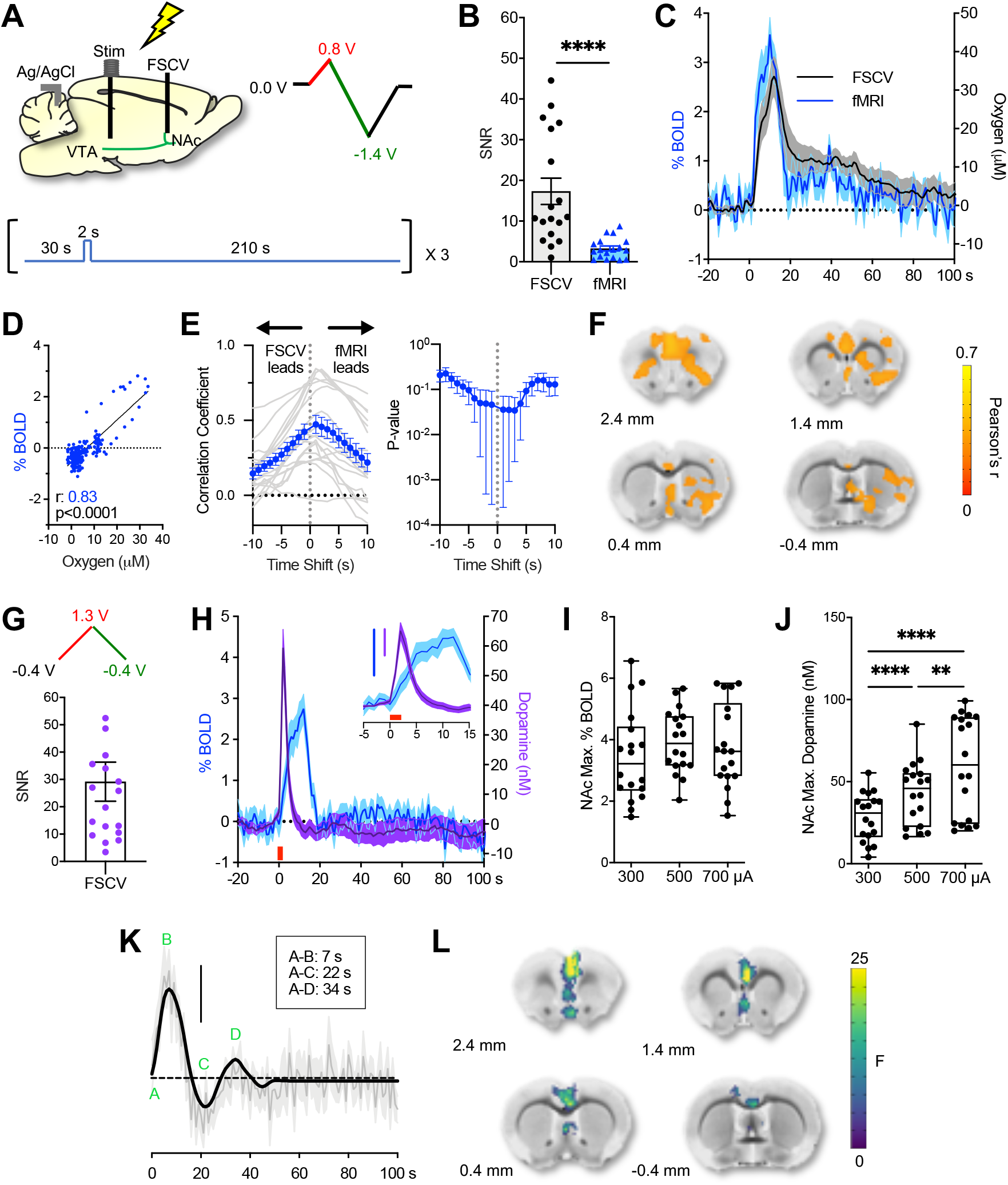
Simultaneous FSCV-fMRI recordings during electrical VTA deep-brain stimulation, using both tissue oxygen FSCV and dopamine FSCV waveforms (2 s, 700 µA, 60 Hz, 2 ms pulse width). (A) Paradigm schematic for electrical stimulations. *Top:* Cartoon of implant locations (*left*), and the oxygen-sensitive FSCV waveform used (*right*). Red indicates the oxidation potential sweep, green indicates the reduction potential sweep. *Bottom*: Electrical stimulation paradigm. (B) SNR values from FSCV and fMRI ROIs during individual stimulation time-courses (n=18 events, ROIs=36±3 voxels; ratio-paired t-test ****p<0.001). (C) Time-courses from NAc recording sites during ipsilateral electrical VTA stimulation (n=18 events). (D) Pearson’s correlation of averaged BOLD fMRI and FSCV oxygen time-courses from c. (E) BOLD fMRI and oxygen FSCV time shift correlations (n=18 events). Individual event time shifts are grey, averaged responses are blue. (F) Group-level correlation maps between FSCV oxygen and BOLD fMRI time-courses (p=0.05). Voxels with correlation coefficients <0.3 are excluded. (G) Dopamine-sensitive FSCV waveform used for the second set of electrical stimulations (*top*), and the SNR for FSCV dopamine (*bottom*; n=18 events). (H) Time-courses from NAc recording sites during the second set of ipsilateral electrical VTA stimulations. Stimulation duration is indicated with a red bar. Inset is a magnification; blue scale bar=2% BOLD response, purple scale bar=20 nM dopamine. (I) Maximum evoked NAc BOLD fMRI values and (J) FSCV dopamine release from different amplitude stimuli (one-way RM-ANOVA, Tukey’s post-hoc **p<0.01, ****p<0.001). (K) Group-level dopamine-derived HRF. Scale bar=2% of BOLD response. Averaged HRF is grey, with smoothed HRF overlaid in black. (L) Linear mixed-effects model maps derived from BOLD fMRI using the dopamine-derived HRF. Group-level main effect of maximum evoked dopamine amplitude (one-way ANOVA, p<0.05 threshold). Error bars are ±SEM.

Next, we recorded and analyzed simultaneous dopamine and BOLD data using FSCV-fMRI. The dopamine-sensitive FSCV waveform recorded high SNR dopamine measurements during fMRI (29.2±7.1; Fig.4G). Evoked dopamine release preceded the hemodynamic response, in agreement with the literature^24^ (Fig.4H), and fMRI responses were consistent (i.e., fMRI responses during dopamine FSCV resembled those obtained during oxygen FSCV in Fig.4C). One-way repeated-measures (RM) analysis of variance (ANOVA) revealed that the maximum evoked BOLD fMRI in the NAc did not significantly change with stimulation amplitude (Fig.4I), but evoked dopamine release did (F_(1.3,22.3)_=39.2, p<0.0001; Tukey’s post-hoc, 300 vs 500 and 700 µA, p<0.0001; 500 vs 700 µA, p=0.0014; Fig.4J). Deconvolution analysis was performed for each simultaneously acquired pair of BOLD and dopamine time-courses to derive subject-specific HRFs (Supplementary Fig.6). The average of all subject-specific HRFs has an initial rise peaking at 7 s, undershoot peaking at 22 s, and a second increase above baseline peaking at 34 s (Fig.4K). Because graded stimulation increased maximum dopamine release but not BOLD amplitudes in the NAc, we used this unique FSCV-fMRI dataset to identify spatial locations where the BOLD response covaried with the released dopamine amplitude. Each dopamine time-course (n=18 events) was convolved with our derived HRF to form a GLM, then used to estimate subject-level contrast maps. The main effect of peak dopamine amplitude on the BOLD response identified significant BOLD activations in the prefrontal cortex and septum, where dopamine neurons also project^38^, as having encoded dopamine amplitude dependence (p<0.05; Fig.4L). Building upon the traditional neuroimaging analysis pipeline, neurotransmitter data from FSCV can provide an additional ground-truth regressor for fMRI data that is crucial for understanding how local neurotransmission can impact brain network dynamics.

## Discussion

FSCV is a minimally invasive electrochemical technique used to detect neurotransmitters such as dopamine at a carbon-fiber microelectrode, and fMRI is a noninvasive way of mapping hemodynamic changes across the entire brain. A platform capable of monitoring dopamine and fMRI simultaneously has been desirable because how dopamine affects brain-wide activity dynamics is poorly understood^22^ and advancing this knowledge could make a tremendous impact on understanding dopaminergic drug mechanisms^22^, human behaviors^19^, and how to improve deep-brain stimulation therapies^21^. Existing FSCV and fMRI studies have used these techniques separately^7,19,20,39^, but one study indicated that simultaneous recordings could be possible^40^. In this study, we develop and characterize simultaneous FSCV-fMRI for the first time, *in vitro* and *in vivo*, within a 9.4 T MRI. Highly resolved oxygen changes via FSCV and concurrent BOLD fMRI detected complementary information about hemodynamics across multiple spatial and temporal scales in response to electrical stimulation and oxygen inhalation challenges (i.e., with and without induced neuronal activity increases). Electrically stimulated dopamine release and simultaneous BOLD fMRI data were collected, and we propose a way to use dopamine time-course information to identify specific brain regions that encode dopamine amplitudes. This technique bridges a critical gap between neurotransmitter activity and neuroimaging data and can help isolate the effect of chemical signaling on brain-wide hemodynamics.

We took advantage of the oxygen-sensing capabilities of FSCV and fMRI to cross-validate evoked hemodynamic changes at multiple scales. The hemodynamic response lags increased neuronal activity by up to several seconds^2^, so it can be observed using conventional FSCV and fMRI sampling rates (10 and 1 Hz, respectively). The spatial resolutions of FSCV and fMRI, however, differ by orders of magnitude (10^5^ µm^3^ and 10^8^ µm^3^ for a 100 µm FSCV microelectrode sensing volume and single fMRI voxel, respectively)^25^. This work required voxel clusters of 36±3 per ROI to obtain detectable fMRI responses, which further increased the spatial mismatch. We observed strong correlations between FSCV and fMRI time-courses in the NAc following oxygen challenges (Fig.3E), in agreement with the only known study to perform simultaneous amperometric tissue oxygen detection and BOLD fMRI^35^; however, FSCV oxygen cleared faster (Fig.3F). Extended hyperoxia decreases venous deoxyhemoglobin percentages, slows cerebral blood flow^41^, and pushes excess dissolved oxygen into surrounding tissues^35^. BOLD fMRI is weighted towards deoxyhemoglobin in venous blood vessels^2^, and FSCV detects tissue oxygen (i.e., oxygen diffused from blood vessels), so these techniques are sensitive to different oxygen dynamics. Thus, slower blood flow extends oxygenated hemoglobin clearance time from vessels within the fMRI ROI relative to tissue oxygen clearance from the much smaller FSCV sampling volume. Despite FSCV’s faster temporal resolution, the onset kinetics of electrically evoked FSCV oxygen responses slightly lagged those of fMRI (Fig.4C,H), in agreement with literature that performed these measurements separately^39^. These data indicate that with respect to tissue oxygen, hemoglobin oxygenation changes more rapidly in response to evoked neuronal activity than oxygen inhalation. It is possible that processes downstream of stimulated neuronal activity may more stringently regulate oxygen delivery within highly localized environments. Combining FSCV with more advanced, vessel-resolution fMRI techniques^42^, or using FSCV arrays with fMRI, could further connect the knowledge gap between hemodynamic responses observed at these two different spatial scales.

Our simultaneous FSCV-fMRI experiments were performed in the NAc because the striatum has the highest concentration of dopamine terminals, and dopamine is a known vasomodulatory neurotransmitter^4,6^. Though the oxygen-sensitive waveform can detect dopamine^24,26^ (Supplemental Fig.1), we quantified dopamine release using data from the dopamine-sensitive waveform^12^ (Fig.4G). The dopamine oxidation potential was shifted anodically during the scan versus during implantation (Supplementary Fig.7), an indication of reference and/or recording electrode biofouling seen with chronically implanted glass carbon-fiber microelectrodes^43^. The shift was not related to the magnetic field or active gradients, as it did not occur *in vitro* (Fig.2B) and did occur in Faraday cage recordings from a subject given the same surgery and recovery period (Supplementary Fig.7). Though post-implantation electrodes showed decreased sensitivity in our hands, others saw consistent longitudinal responses^30^. Strategies to mitigate chronic implantation sensitivity problems are being researched^44,45^, but our data indicate that reliable, high SNR analyte detection during fMRI can still be achieved days after electrode implantation.

The ability to observe dopamine dynamics and whole-brain imaging simultaneously would enormously benefit research into global brain circuitry changes from drug abuse, pain, or disorders implicated in dopamine dysfunction^7,10,19,22^. By measuring and quantifying evoked dopamine release, FSCV-fMRI can generate data-driven HRFs (Fig.4K) and help identify brain regions sensitive to different dopamine release amplitudes (Fig.4L). These data can be used to study effects of neuronal or pharmacological manipulations and tease apart the role of dopaminergic signaling on local and global hemodynamic responses in healthy and pathological brains^10,22^. Our study chose an electrical stimulation protocol for FSCV to induce reliable dopamine changes in the NAc^24^; however, neurochemicals other than dopamine are released in the NAc following electrical or optogenetic VTA dopaminergic neuron stimulation^46^, so these results do not establish dopamine as the sole neurochemical source responsible for the observed oxygen changes. Future studies that selectively drive dopamine release with confounding factors eliminated (e.g., suppressing neuronal activity and/or unwanted neurotransmission) will further isolate the role of dopamine and its receptors^7,36^.

Though our electrical stimulations lack specificity, the analysis framework proposed herein should set the foundation for neurotransmitter-based HRF modeling in the future. Interestingly, our dopamine-modeled HRF has two notable peaks, different from the single-peak canonical HRF^47^. The first peak is the initial rise, and an additional peak occurs after the first post-stimulus undershoot. This “second peak” is well documented in the oxygen FSCV literature using both VTA and median forebrain bundle stimulations^24,26,27^, supporting our approach towards deriving an HRF to describe the relationship between concurrently measured neurochemical release and BOLD fMRI. Literature attributes these oxygen increases to cerebral blood flow^27^, and that each peak is mediated differently; for example, an adenosine receptor antagonist decimated the second peak response but had no effect on the first^24^. Interestingly, blocking dopamine synthesis had no effect on the either oxygen peak^27^. More research is needed to isolate dopamine and dopamine receptors’ influence over HRFs.

Our method using a ground-truth output metric (i.e., quantified dopamine release amplitudes) lends additional neurochemical context to response map interpretations and can pull out relationships that traditional input models (e.g., stimulation paradigms) would miss. For instance, though a traditional stimulus paradigm model with GLM pulled wide-spread activation around ipsilateral NAc (Supplementary Fig.5) similar to literature^7,36,37^, maps of dopamine amplitude as the main effect isolated prefrontal cortex and septum (Fig.4L). Our method utilizing ground truth data to guide models is more effective than relying on assumptions that the evoked response would scale with the given stimulus inputs, and can account for inter-subject response variability. The ability to identify locations where brain activity co-varies with quantitative neurotransmitter amplitudes could have far-reaching benefits for research studying local neurotransmission influence over brain hemodynamics and behavior.

Though FSCV has been a common method of detecting *in vivo* neurotransmitters for decades, an increasing number of optical detection alternatives exist that could also pair with fMRI^17,48–50^. A dopamine-sensitive protein-based MRI contrast agent has been used with BOLD^13^, and dopamine dynamics have been explored using near-infrared fluorescent sensors^15^ and genetically encoded fluorescent sensors^14,16^. These techniques require either infusion cannulae or optical fibers for contrast agent dispersal or fluorescence detection, respectively, which use fMRI-compatible materials. These alternative dopamine detection methods have high spatiotemporal resolution, even exceeding FSCV, making them exciting potential multimodality expansions. These methods show qualitative percent changes in signal, so quantitative FSCV measurements could provide useful cross-validation. FSCV-fMRI remains a strong multimodality for studying dopaminergic influence on neurovascular coupling, as it requires a single implantation surgery, no genetic modifications or injections, and is the only current method available to measure electroactive neurotransmitter activity and whole-brain hemodynamics simultaneously. Importantly, FSCV is the fastest-resolution technique to quantify neurotransmitter release in humans^12,18–20^, which is crucial for detecting rapid neurotransmission events during behavior. Expanding this multimodality to include other neurotransmitter sensors or other techniques is a realistic possibility that could hone in on the role of dopamine in the hemodynamic response at multiple spatiotemporal scales in both preclinical^22^ and clinical^21^ experiments.

## Materials and Methods

### Fast Scan Cyclic Voltammetry (FSCV)

Carbon-fiber microelectrodes were fabricated as previously described^30^, with modifications for MRI compatibility. Briefly, a 5-7 µm-diameter carbon fiber was manually threaded through a saline-filled, polyimide-fused silica capillary. The capillary was sealed with clear epoxy at the recording end such that 100-200 µm of the fiber was left exposed, and the other end of the fiber was sealed to a silver wire with silver epoxy and silver paint. The silver wire/carbon fiber junction was encapsulated with clear epoxy for stability.

FSCV acquisition hardware was custom-built within MR-compatible aluminum casing (UNC Electronics Facility, Chapel Hill, NC, USA), and data were collected and analyzed with the High Definition Cyclic Voltammetry (HDCV) computer program^51^. The standard coaxial cables used to carry signal from the electrodes to the headstage were replaced with MR-compatible triaxial cables. Triaxial cables were chosen to prevent current leakage via a driven shield, as well as to compensate for the additional cable capacitance introduced by the length of the cables necessary to connect the headstage to the electrodes through the back of the MRI bore (approximately 2 m). Output digitization was smoothed by passing digital waveform signals through a 2 kHz low-pass analog filter before being applied to the electrode. Input signals were split via T adapter and collected as raw current at one input channel (for analysis) and passed through an analog 2 kHz low-pass filter prior to collection on a second input channel (for real-time signal viewing). Ferrites were placed on both input and output voltage coaxial and triaxial cables to further eliminate high-frequency noise (Fair-Rite Products Corp., Wallkill, NY, USA). A 4^th^ order Bessel low-pass filter (2 kHz cutoff frequency) was digitally applied to input currents post-acquisition.

Waveforms were applied to the microelectrodes to selectively oxidize and/or reduce dopamine and/or oxygen (Supplementary Fig.2). The oxygen-sensing waveform is an 11 ms waveform that holds at 0 V, scans up to +0.8 V, down to –1.4 V, then back to 0 V^24–26^. The dopamine-sensitive waveform is an 8.5 ms, triangular waveform with a −0.4 V holding potential that scans to +1.3 V and back to −0.4 V^24,52^. Waveforms were applied at 5 or 10 Hz, voltages were scanned at 400 V/s, and data were sampled at 100 kHz. All voltages are versus Ag/AgCl. In both calibration and *in vivo* studies, the oxygen-sensitive waveform was used prior to the dopamine-sensitive waveform.

A “dummy cell” was fabricated to characterize noise within the MRI bore, which consisted of a 30 kΩ resistor and 0.33 nF capacitor connected in series on a circuit board, to mimic solution resistance and carbon fiber electrode double-layer capacitance, respectively. Dopamine stock solutions and oxygen calibration solutions were freshly prepared in room temperature phosphate-buffered saline (pH 7.4). Flow-injection analysis was used to collect the current amplitude responses to an analyte bolus of known concentration^53^. Concentrations from 100-2000 nM were injected in random order, and each concentration was injected in triplicate. The within-bore MRI flow-injection datasets were acquired with the gradient disabled with low-pass filtering, enabled without low-pass filtering, and enabled with low-pass filtering. These datasets were acquired in random order to ensure that noise from the enabled gradient had no detrimental effect on electrode sensitivity. Flow injection analysis data from within the MRI bore were not used to calibrate electrodes, as the tubing length required to keep the magnetic 6-port injector a safe distance from the MR heavily diluted the bolus concentrations. Flow-through analysis was repeated inside a Faraday cage to obtain calibration factors for pre- and post-implantation electrodes. Only oxygen waveform calibration data where the oxygen waveform was applied first are included (i.e., those that used the dopamine waveform first were excluded).

### Animal Care

All animal protocols were approved by the Institutional Animal Care and Use Committee of the University of North Carolina at Chapel Hill (UNC) in accordance with the Guide for Care and Use of Laboratory Animals (eighth edition). Nine male Long-Evans rats (300–450 g, Charles River, Wilmington, MA, USA) were pair-housed prior to surgery at UNC animal facilities, given food and water ad libitum, and kept on a 12 h light/dark cycle. Care was taken to reduce the number of animals used. Three animals were excluded: Two animals had prohibitively large imaging artifacts, likely from the connective leads plugged into the stimulating electrode pedestal in the slices of interest, and one animal’s electrode stopped functioning before the experiment finished.

### Surgery

Rats were deeply anaesthetized with 4% isoflurane, placed in a rodent stereotaxic frame (Kopf, Tujunga, CA, USA), and maintained at 2% isoflurane. Depth of anesthesia was confirmed via toe pinch and eye blink responses. Coordinates are relative to skull at bregma^54^. Holes were drilled over the ventral striatum (1.3 mm A-P, 2.0 mm M-L, 6.8-7.8 mm D-V) for the carbon-fiber electrode and the VTA (−5.2 mm A-P, +1.2 mm M-L, −8.2 to −9.2 mm D-V) for the bipolar, MR-compatible twisted tungsten stimulating electrode (PlasticsOne, VA, USA); to avoid the sagittal sinus, the stimulating electrode was implanted at a 4 degree angle. The stimulating electrode was untwisted enough to separate the tips by 1 mm^55^. The D-V coordinates of the stimulating and working electrodes were optimized to evoke maximal dopamine release. The dopamine-sensitive waveform was used during implantation, so oxygen responses to stimulation were unknown prior to fMRI scanning. An additional 3-4 burr holes were drilled for MR-compatible, brass skull screws. An Ag/AgCl reference electrode wire was placed in the contralateral hemisphere. Silicone adhesive was applied around the working electrode to prevent cortical tissue damage from the heat of curing dental cement (World Precision Instruments, Sarasota, FL, USA). Skull screws, electrodes, and plastic shield were held in place using dental cement. Subjects were singly-housed for at least 4 days to recover before scanning.

### Phantom MRI

All MRI data were acquired using a Bruker 9.4 T MRI. *In vitro* scans used a Bruker 72 mm volume coil. An FSCV electrode was embedded within an agarose phantom and scanned to examine whether the materials induced artifacts in RARE (BW=312.5 kHz, TR=2000 ms, TE=40 ms, in-plane resolution= 100 µm, slice thickness=100 µm) or FLASH (BW=312.5 kHz, TR=50 ms, TE=3.52 ms, in-plane resolution= 100 µm, slice thickness=100 µm).

### FSCV-fMRI in vivo

*In vivo* BOLD fMRI data were acquired using a homemade surface coil, and subjects were prepared as described previously^36,56^. Briefly, subjects were anesthetized and paralyzed by infusing a cocktail of dexmedetomidine and pancuronium bromide (0.1 mg/mL and 1.0 mg/mL respectively, intraperitoneal) to prevent motion artifacts, supplemented with low dose isoflurane (0.5%), and artificially ventilated via endotracheal intubation. Experiments did not begin until rectal temperature stabilized at 37±0.5 °C, end-tidal CO_2_ was 2.6-3.2%, and oxygen saturation was ≥95%. Physiology was monitored throughout the experiment and maintained within normal ranges. Five-slice echo planar imaging (EPI) data were acquired with BW=333.33 kHz, TR=1000 ms, TE=15 ms, 80×80 matrix, FOV=2.56 cm^2^, slice thickness=1 mm. One slice was discarded prior to data analysis due to artifacts from the stimulating electrode. EPI slice acquisition and electrical stimulations were both controlled via HDCV software. To interleave fMRI and FSCV data acquisition, HDCV sent a 10 ms, high-to-low TTL pulse to trigger EPI per-slice acquisition 3 ms after each applied FSCV waveform.

Oxygen challenges were administered and repeated in triplicate, where 100% medical air was inhaled for 60 s, then 100% oxygen for 60 s, and finally 100% medical air for 180 s. For deep brain stimulations, biphasic square-wave stimulations were administered 180 s apart, at 60 Hz with current amplitudes of 300, 500, and 700 µA, a 2 ms pulse-width, and total duration of 2 s. Each stimulation amplitude was administered in triplicate and in random order with simultaneously acquired oxygen FSCV, followed by a repeated set of randomly ordered triplicate stimulations per amplitude with simultaneously acquired dopamine FSCV. Finally, a powerful stimulation was given to evoke large responses for principal component analysis (PCA) training sets needed for FSCV analysis (60 Hz, 900 µA, 4 ms pulse-width, 4 s duration). A high-resolution RARE scan was obtained at the end of each scan to verify each electrode location (BW=183.1 kHz, TR=2500 ms, TE=33 ms, in-plane resolution=100 µm, slice thickness=1 mm).

### Data Analysis

FSCV data were analyzed within the HDCV analysis software, which can be licensed to academic users through the University of North Carolina Electronics Facility (University of North Carolina at Chapel Hill, Chapel Hill, NC, USA). FSCV data were background subtracted and filtered with a 4^th^ order, 2 kHz cutoff low-pass Bessel filter. FSCV background currents were taken prior to each gas challenge or stimulation event. Post-implantation calibration factors from flow-through analysis, normalized to the length of the exposed carbon fiber, were used to quantify oxygen and dopamine concentrations. To verify that all faradaic currents resulted from oxygen or dopamine, data sets were analyzed using principal component analysis (PCA) within the HDCV analysis software^51^. Training sets for PCA consisted of 4-8 different concentrations per analyte of interest, per electrode. FSCV SNRs were calculated by subtracting a 10 s baseline average prior to stimulation from the maximum peak oxidative (dopamine) or reductive (oxygen) current, then dividing the difference by the standard deviation of the same 10 s baseline period. FSCV data were binned to 1 s increments for all fMRI time-course comparisons.

BOLD fMRI data were pre-processed and analyzed in Analysis of Functional NeuroImages (AFNI) software (National Institute of Mental Health, Bethesda, MD, USA). Prior to preprocessing, all fMRI data were zero-padded to ensure that the data had a consistent and sufficient matrix size to be used as inputs for the AFNI image registration software. T2 anatomical and EPI fMRI data were manually aligned to a template using ITK-Snap. The first 8 TRs were excluded to account for steady state, and the first 8 s of FSCV time-courses were discarded to match. Data were interpolated in order to follow AFNI pre-processing pipeline slice requirements. After pre-processing, data matrices were truncated to their original size. BOLD time-courses were extracted using an ROI mask unique to each subject, based on the location of the tip of the FSCV microelectrode (Supplementary Fig.4).

Head motion and baseline drift were removed by linear regression prior to first-level statistical analyses and ROI time-course extractions. Subject-level correlation maps were calculated using BOLD fMRI and oxygen FSCV time-courses from oxygen challenge and electrical stimulation experiments. Fisher z-transformations were performed on subject-level oxygen challenge and electrical stimulation correlation maps for group statistics. A one-sample t-test was performed using linear regression to identify voxel clusters that fit with the intercept line representing the average, at p<0.05. Benjamini/Hochberg false discovery rate (FDR) correction was used to correct the voxel-level multiple comparison problem, with a q<0.05 threshold. Then, a group average correlation map was generated using the surviving clusters. Subsequently, the inverse Fisher’s Z-transform was performed to restore Fisher Z to Pearson’s R correlation coefficients.

Traditional functional activation maps were obtained to compare with our hemodynamic response function (HRF) derived maps. The BOLD response was first predicted by convolving the electrical stimulus (i.e., 2 sec block) and the following gamma function (i.e., the default HRF in AFNI): HRF(t) = int(g(t-s), s=0..min(t,d)), where g(t) = t^q*exp(-t) /(q^q*exp(-q)), where t=time and q=5. This stimulus-derived HRF predictor was used as a regressor for subject-level GLMs to identify significantly activated voxels from individual response maps. Group-level analyses used a one-tailed t-test to test mean subject-level regression coefficient maps against zero, with a q=0.05 FDR threshold.

Oxygen ROI time-courses were low-pass filtered with a 0.05 Hz cutoff to remove physiological noise prior to deriving kinetic parameters (oxygen challenge) and subject-level HRFs (electrical stimulation). The HRF was derived using the linear system: BOLD(t) = DA(t) * HRF + baseline + linear drift(t), where DA is dopamine. DC offset and linear drift were removed from BOLD time-courses during HRF deconvolution analysis (Supplementary Fig.6). The averaged HRF was smoothed with a 3^rd^ order Savitzky Golay filter. To identify brain regions that encode dopamine or stimulation amplitude, the smoothed HRF was convolved with dopamine time-courses to form Bayesian Ridge GLM regressors, which were subsequently used to estimate subject-level BOLD fMRI contrast maps. Second-order polynomial curves were added to detrend the BOLD signal prior to statistical group-level analysis.

## Code Availability

Standard AFNI codes were used for BOLD fMRI pre-processing and traditional functional activation map analysis. MATLAB (The Mathworks, Inc., Natick, MA, USA) codes for the time shift analysis and HRF convolution will be made available upon request, as will Python codes used to obtain statistical response maps and correlation maps.

## Statistical Information

Statistics were performed using GraphPad Prism 8 (GraphPad Software, San Diego, CA, USA), AFNI, and Python. RM-ANOVA was used to compare *in vitro* SNRs between multiple electrodes, with gradient and filter status as the within-subject factor. Maximum evoked BOLD and dopamine RM-ANOVA analyses were performed using stimulation amplitude as the within-subject factor. Significant ANOVA interactions were subjected to Tukey’s post-hoc multiple comparisons analysis. FSCV and fMRI values for *in vivo* SNR data and oxygen challenge kinetic parameters were compared using two-tailed, ratio-paired t-tests.

One-way ANOVAs were used to test the main effects of the maximum evoked dopamine amplitude on HRF-derived contrast maps, using a permutation method (5000 times iteration) with the maximum T method for multi-voxel, multiple-comparison correction. A p<0.05 threshold was used to indicate statistical significance in all analyses, and all statistical clusters were limited to clusters >50 voxels to exclude potential false positives.

## Acknowledgements

We acknowledge the help of Dr. Weiting Zhang for her analysis code, Dr. Domenic Cerri for helpful discussions, and members of the UNC Center for Animal MRI (CAMRI) and Biomedical Research Imaging Center (BRIC) for technical assistance. Research reported in this publication was supported by the National Institutes of Health under Award Number F32MH115439, RF1MH117053, R01NS091236, R01MH111429, R41MH113252, P60AA011605, U01AA020023, R01AA025582, and P50HD103573. The content is solely the responsibility of the authors and does not necessarily represent the official views of the National Institutes of Health.

## Author Contributions

L.W. designed and ran all experiments, preprocessed and analyzed data, and prepared the manuscript. M.V. helped characterize the multimodality and troubleshoot electronic problems, and fabricated custom hardware. S.L. performed the majority of the fMRI statistical analyses, and, along with T.C., assisted in the design, execution, and interpretation of the HRF analysis. L.W., R.M.W., and Y.Y.S. conceived the research idea. R.M.W. and Y.Y.S. provided advice on experimental designs throughout the project. All authors have read, commented on, and approved this manuscript.

