## Supplemental Figures for "Simultaneous fMRI and fast-scan cyclic voltammetry bridges oxygenation and neurotransmitter dynamics across spatiotemporal scales"

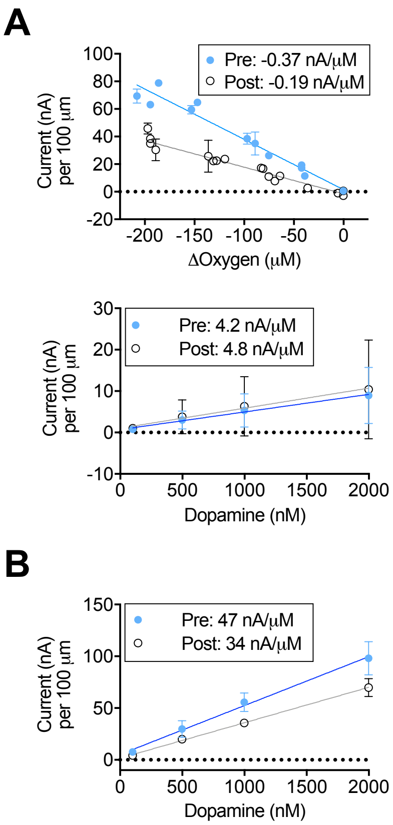


**Supplemental Fig. 1** Calibration constants derived from flow-through analysis at pre- and post-implantation polyimide-fused silica carbon-fiber microelectrodes. (A) Sensitivity for oxygen (top; n=3 pre, n=5 post) and dopamine (bottom; n=7 pre, n=5 post) on the oxygen-sensitive waveform. ΔOxygen indicates the difference in oxygen concentration versus air-saturated buffer at 22°C (225 μM). (B) Sensitivity to dopamine is greater using the dopamine waveform (n=5 pre, n=3 post). Values are triplicate measures per electrode shown as averages +/- SD.

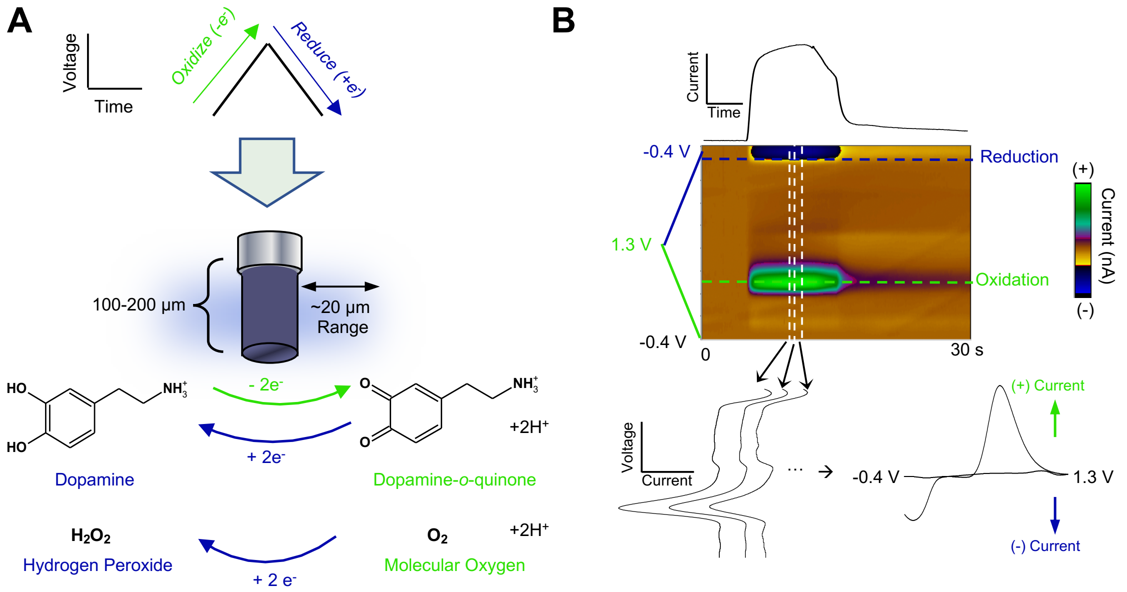


**Supplementary Fig. 2** FSCV detects electrochemically active analytes. (A) Voltage sweeps (waveforms) are applied to a carbon fiber microelectrode at a specified frequency and scan rate to electrochemically oxidize and/or reduce chemical analytes (i.e., subtract and/or add electrons [$e^-$], respectively). (B) FSCV data is displayed in 3D color plots. *Top*: A dopamine time-course from the oxidation potential of dopamine (horizontal green dashed line). *Middle*: Color plot of a dopamine bolus, where time is on the x-axis, the applied voltage is on the y-axis, and current amplitudes are indicated in false color. *Bottom*: Vertical slices give voltage versus current plots at a specified time-point (vertical white dashed lines). Because waveforms cycle through the same voltages, these can be folded into a cyclic voltammogram that visually indicates the chemical(s) detected at the microelectrode surface based on oxidation and reduction potential peaks on a plot of voltage (x-axis) versus current amplitude (y-axis).

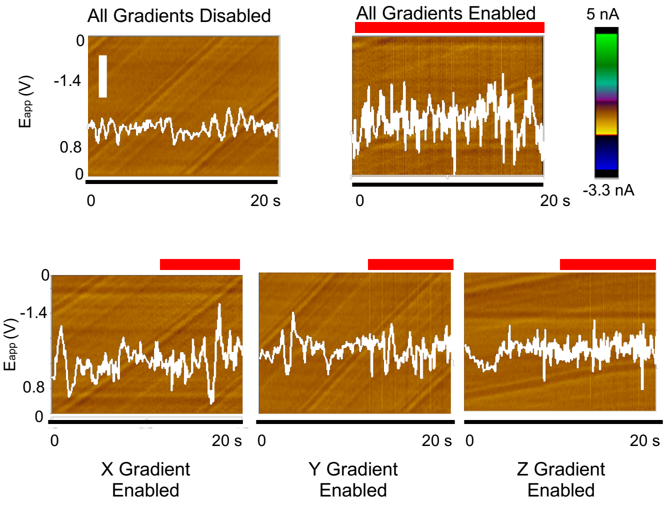


**Supplementary Fig. 3** Low-pass filtered (2 kHz cutoff), baseline electrical noise at a dummy cell within a 9.4 T MRI bore when selected gradients are enabled, but without active pulsing. Current time-courses taken from –1.0 V are overlaid in white, and red bars indicate the time of individual gradients being enabled. Vertical white scale bar=0.2 nA.

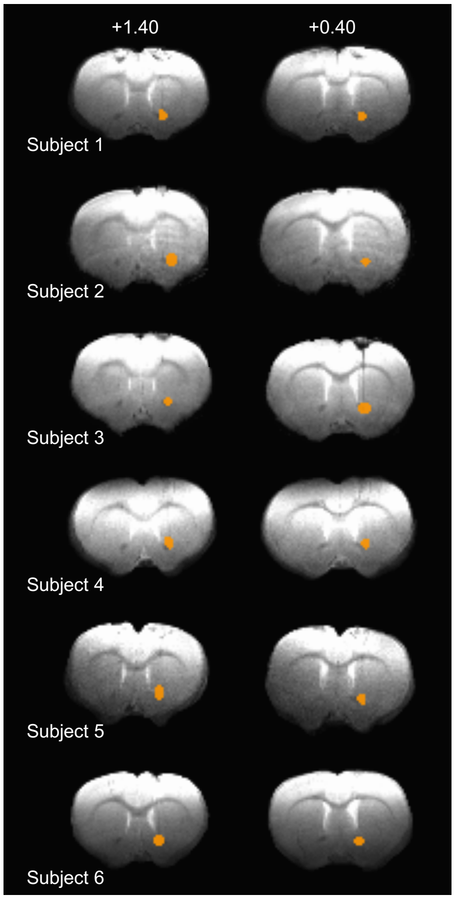


**Supplementary Fig. 4** Histological verification of electrode locations. Electrode-focused ROIs drawn for each subject based on electrode tip location, in orange. The ROI includes multiple voxels (average size=36±3 voxels) to account for the low SNR in BOLD fMRI.

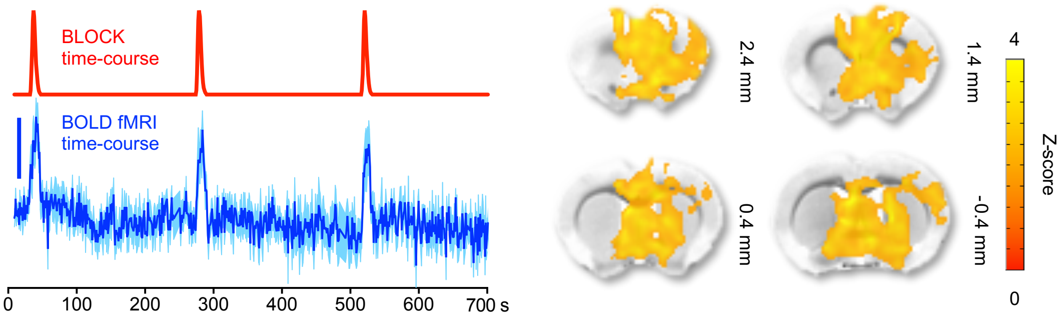


**Supplementary Fig. 5** Electrical VTA deep-brain stimulation data from Fig.4H analyzed using a stimulus paradigm-based fMRI analysis. A block design paradigm based on the stimulation duration (left; 2 s stimulations) was convolved with a canonical HRF for GLM analysis (right, one-sided t-test with q=0.05 threshold, n=6 subjects). Error bars are ±SEM. Scale bar for BOLD fMRI time-course=2% signal change.

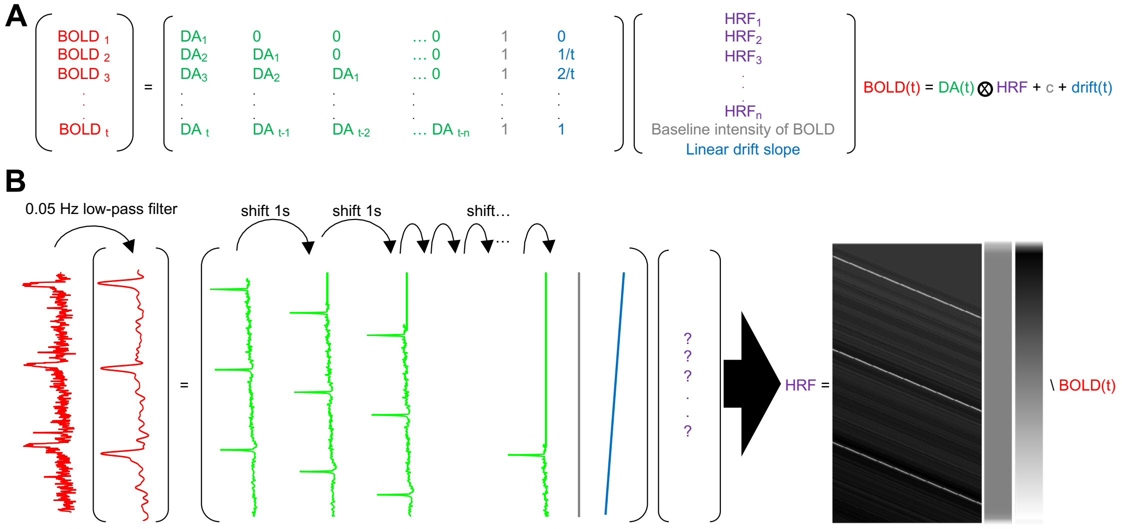


**Supplementary Fig. 6** Deriving an HRF from neuronal and dopaminergic information. (A) Deconvolution matrices are used to derive HRFs from concurrently collected BOLD fMRI and dopamine time-courses. DA=dopamine. $BOLD_1$ indicates the BOLD fMRI time-course value at time (t)=1, $BOLD_2$ is at (t)=2, etc. (B) Representative data showing the HRF deconvolution process.

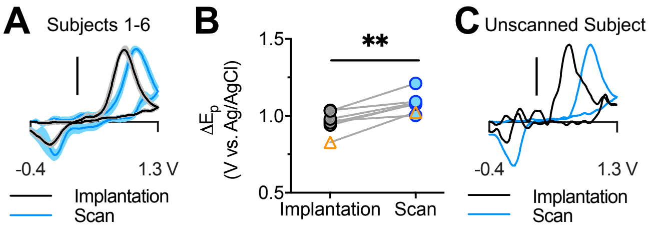


**Supplementary Fig. 7** Implantation time affects dopamine sensitivity *in vivo*. (A) Evoked dopamine release cyclic voltammograms from the day of implantation (black) and in recovered animals during scanning after 4 days of recovery (blue). Averages ±SEM are shown. (B) Quantified changes in dopamine cyclic voltammogram peak separations. Data obtained in a Faraday cage from (C), a subject given the same surgical procedures and recovery time as scanned animals, are shown as open orange triangles. All voltammograms are normalized so that the maximum oxidation current equals 1. All scale bars indicate 0.5 of maximum normalized value. Paired t-test **p<0.01.
